# Benchmarking DNA isolation kits used in analyses of the urinary microbiome

**DOI:** 10.1101/2020.11.10.375279

**Authors:** Lisa Karstens, Nazema Y. Siddiqui, Tamara Zaza, Alecsander Barstad, Cindy L. Amundsen, Tatyana A. Sysoeva

## Abstract

The urinary microbiome has been increasingly characterized using next-generation sequencing. However, many of the technical methods have not yet been specifically optimized for urine. We sought to compare the performance of several DNA isolation kits used in urinary microbiome studies. A total of 11 voided urine samples and one buffer control were divided into 5 equal aliquots and processed in parallel using five commercial DNA isolation kits. DNA was quantified and the V4 segment of the 16S rRNA gene was sequenced. Data were processed to identify the microbial composition and to assess alpha and beta diversity of the samples. Tested DNA isolation kits result in significantly different DNA yields from urine samples but non-significant differences in the number of reads recovered, alpha, or beta diversity. DNA extracted with the Qiagen Biostic Bacteremia and DNeasy Blood & Tissue kits showed the fewest technical issues in downstream analyses, with the DNeasy Blood & Tissue kit also demonstrating the highest DNA yield. The Promega kit recovered fewer Gram positive bacteria compared to other kits. The Promega and DNeasy PowerSoil kits also appear to have some important biases towards over-representing certain Gram negative bacteria of biologic relevance within the urinary microbiome.

## Introduction

Resident microbes in multiple niches of the human body are being studied for their impact on health and disease. The microbiome of the urinary tract has not been extensively characterized though differences in urinary microbiota are evident in urologic conditions such as urgency urinary incontinence ^1–3^. It is also likely that the presence of urinary commensals affects the propensity towards development of urinary tract infections^4^. It is now known that urine contains a range of fastidious bacteria that are not detected using standard urine culture, or even with the recently developed enhanced quantitative urine culture (EQUC) techniques ^1,5^. As such, characterization of the urinary microbiome has been accomplished using culture-independent methods, relying on next generation sequencing methods such as bacterial 16S rRNA gene sequencing, also known as amplicon sequencing or marker gene sequencing.

Despite the fact that the urinary microbiome has been recognized for almost a decade,^1^ many of the technical methods used in marker gene sequencing have not been optimized or standardized for detection of urinary microbiota, as has been done for other microbiome niches ^6,7^. When performing DNA sequencing on biological samples to extract information about the bacterial communities present, multiple technical steps are required in order to name and classify the microbes contained in the sample: sample collection, storage and handling, DNA isolation, amplification, and sequencing ^8^. At each of these steps, bias could influence the final results, and several have been evaluated in recent studies. The method of sample collection affects the recovered urinary microbiome characteristics, with catheterized sampling offering the most specificity for the bladder environment^1,9,10^. As for storage and handling steps, high concentrations of certain chemicals in urine result in precipitation of crystalline and amorphous materials such as uric acid, calcium phosphates, calcium oxalate and others ^11^. The presence of crystalline precipitants in urine were recently shown to alter the pelleting and lysis of cells, and biochemical reactions such as amplification via polymerase chain reaction (PCR) prior to sequencing^12,13^. Storage and handling of urine specimens after collection has also been investigated ^14^, and demonstrate that urine samples should be cooled as soon as possible if a stabilizing agent such as Assay Assure^®^ is not used.

In addition to sample collection, handling and storage, the DNA isolation methods used are another important step for microbiome analysis where bias could be introduced prior to sequencing. At this step, human and microbial DNA are extracted from the proteins, salts, and other components of the physiologic sample. This requires lysis of human cells and bacterial cell walls in order to isolate the DNA contained within. When performing marker gene sequencing, the isolated DNA is later subjected to PCR, where the marker gene is amplified and uniquely tagged for sequencing. The bacterial 16S rRNA gene is one of the most commonly used marker genes used for bacterial identification. To date, a range of amplicons encompassing multiple different variable regions of the 16S rRNA gene including V2, V3-V4, V4, V4-V5, and V6 ^8,15,16^ have been applied to urinary microbiome samples.

Regardless of which segment of DNA is used as the marker gene, reliably isolating all of the DNA in a sample is an important step prior to PCR and sequencing. Many commercial kits and custom protocols for DNA isolation were developed for microbiome analyses specifically for microbe-rich or microbe-poor environments. This tailoring of DNA isolation methods was required to achieve more representative identification and quantification of the microbial composition for each respective environment. Nevertheless, different methods for DNA isolation show variable efficiencies of DNA recovery and quality. A large number of studies report significant differences in microbial composition identified with the use of different DNA isolation protocols^17–24^ Biases introduced by the DNA isolation methods to microbial composition persist both in microbe-rich communities such as gut, soil, sewage ^19,25^ and in microbe-poor communities such as water, meconium, and animal larvae ^22,26^. However, there are occasionally studies that do not show notable differences among DNA extraction methods in other microbial niches ^23^.

Most studies that examine differences among DNA extraction protocols note that the main hurdles are incomplete cellular lysis and presence of PCR inhibitors that could interfere with downstream sequencing. Incomplete cellular lysis for some of the microbes biases the compositional analyses towards more easily lysed taxa. These differences in lysis were repeatedly recorded for Gram positive bacteria and fungi that both have more robust cell walls in comparison with the Gram negative bacteria ^20,26^. For the urinary microbiome, researchers already found a significant bias in the ability to detect fungi due to the inability to efficiently lyse hardy fungal cells ^13,27^. However, many urinary microbiome studies employ a variety of DNA isolation techniques without considering these potential sources of bias. Table S1 summarizes these studies to highlight the diversity of the methods of DNA isolation used to date. These studies use both custom and commercially available DNA isolation methods. As there are no studies directly comparing the results obtained when urine samples are subjected to different commercial DNA isolation kits, our primary objective was to assess whether recovered microbes identified by 16S rRNA sequencing differ based on the DNA isolation protocol.

## Results

### DNA Recovery & Performance in High Throughput Sequencing

A total of 11 urine samples and one negative control containing phosphate buffered saline (PBS) were equally divided and subjected to parallel DNA isolation procedures with five DNA isolation kits (Table 1). The total DNA concentration recovered from each DNA isolation kit was highly variable (Figure 1A, Table S3) with the Qiagen DNeasy Blood and Tissue kit resulting in the highest concentrations compared to the others (Kruskal-Wallis p = 0.0007). Since each aliquot from urine samples contained the same starting material per kit, and each kit elutes DNA into the same volume, higher concentrations would reflect a higher total amount of DNA isolated. Of the 60 samples (55 from urine and 5 controls), a total of 7 (11.6%) did not produce identifiable bands on gel electrophoresis after PCR amplification of the V4 region of the 16S rRNA gene (Table S3, Figure S1A). The majority of these samples were derived from one urine specimen with the lowest quantity of recovered DNA (Sample 11) and negative controls, suggesting truly low quantity DNA in these samples. However, in one instance (Sample 4), no gel band was detected after PCR when DNA was extracted with the Qiagen DNeasy Ultraclean kit though bands were identified when DNA was isolated with all of the other four kits.

**Table 1.** Characteristics of microbial DNA isolation kits used in this study.

| Kit | Full name | Mechanical lysis | Enzymatic lysis | Chemical lysis | Heat treatment | Gram+ adaptation | DNA binding |
| --- | --- | --- | --- | --- | --- | --- | --- |
| BiOstic | Qiagen BiOstic Bacteremia | Yes | No | Yes (G) | Yes | No | Silica spin-column |
| Blood& Tissue | Qiagen DNeasy Blood and Tissue | No | Yes (L, PK) | Yes (D, G) | Yes | Yes | Silica spin-column |
| Promega | Promega Maxwell RSC Purefood GMO and Authentication | No | Yes (PK) | Yes (G) | Yes | No | Cellulose-based particles |
| PowerSoil | Qiagen DNease PowerSoil | Yes | No | Yes (G) | No | No | Silica spin-column |
| UltraClean | Qiagen DNeasy UltraClean | Yes | No | Yes (D, G) | Yes | No | Silica spin-column |

\*L – lysozyme, PK – proteinase K, G – guanidine salts, D – detergent

**Figure 1.**
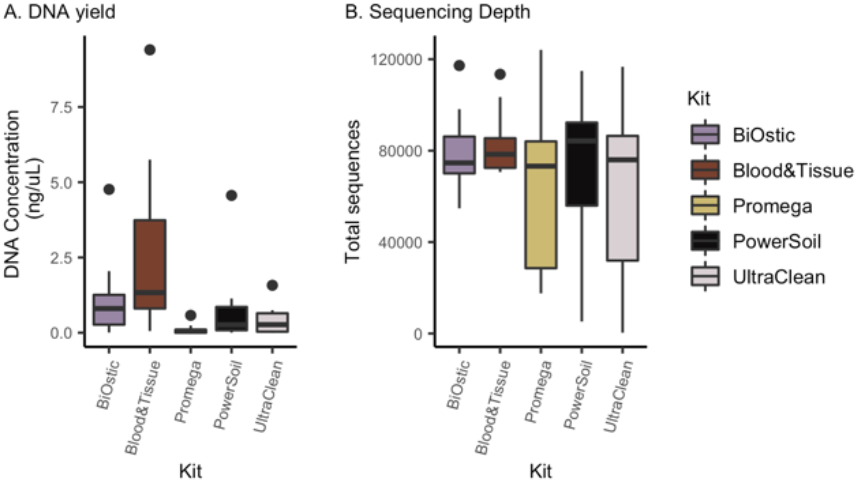
Isolation kits produce different total DNA concentrations but similar 16S specific sequencing depth. **A.** DNA concentration measured by Qubit varied significantly by DNA isolation kit used (Kruskal-Wallis p = 0.0007). **B.** Differences in total DNA concentration did not translate to significant differences in the number of sequence reads per sample (Kruskal-Wallis p = 0.806).

Despite the differences in DNA concentrations between isolation kits, DNA isolated from all kits appeared to perform similarly in high throughput sequencing. We did not identify significant differences in the total number of recovered reads based on the DNA isolation kit (Kruskal-Wallis p = 0.806, Figure 1B). Notably, sequencing reads were obtained even in samples without gel bands after PCR that might have originally been presumed to be devoid of DNA.

### Microbial Composition

Alpha diversity measures summarize the composition of bacteria in a sample in terms of the numbers of different taxa present (richness) and their distribution (evenness). We did not identify significant differences in alpha diversity measured as the number of observed genera, the Shannon index, or the inverse Simpson index based on DNA isolation kit (Kruskal-Wallis p = 0.292, 0.363, and 0.436, respectively; Figure 2).

**Figure 2.**
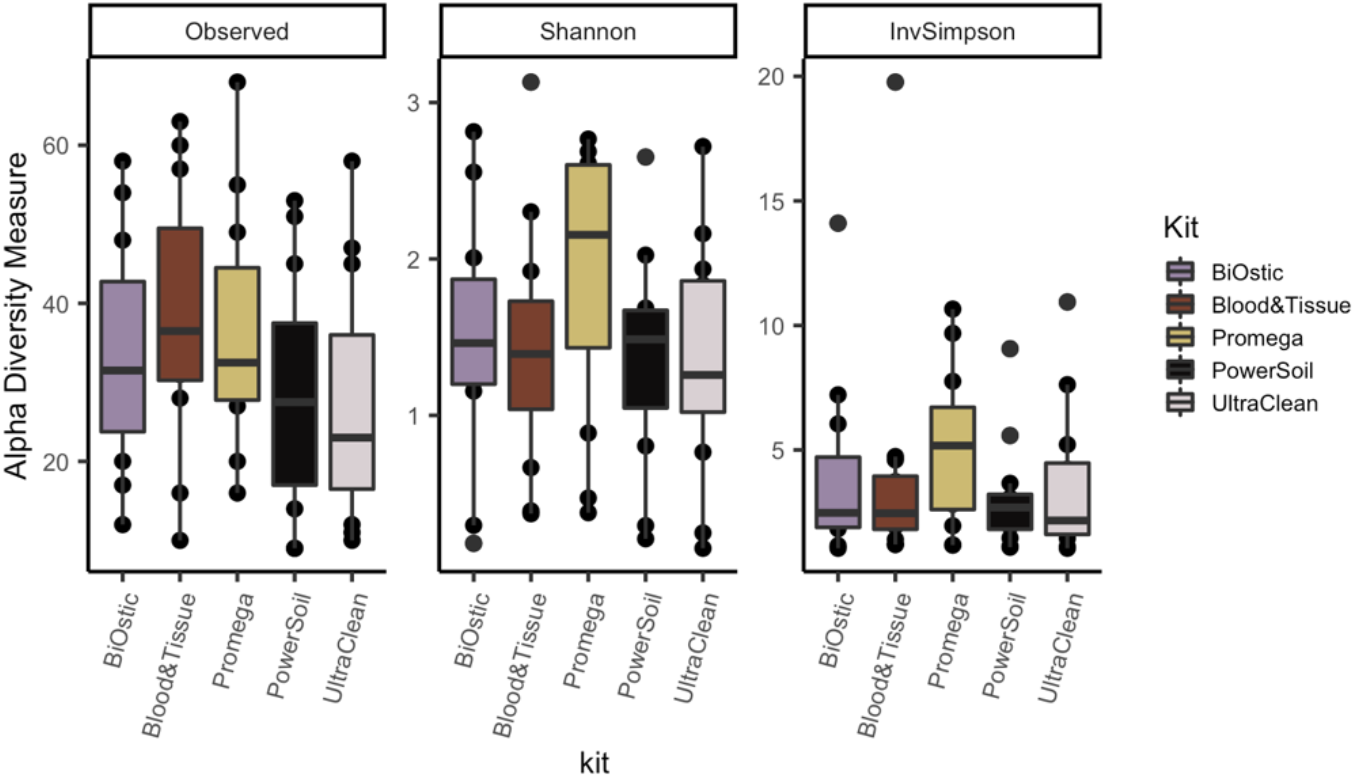
Richness and evenness of the microbial composition does not depend on the testing DNA isolation kit. (Kruskal-Wallis p = 0.292, 0.363, and 0.436, respectively).

To evaluate the differences in the overall composition of taxa between DNA isolation kits, we estimated beta diversity using the Bray-Curtis distance and nonmetric multi-dimensional scaling (NMDS, Figure 3A), and evaluated the relative abundance of recovered bacteria in each sample (Figure 3B). For most of the samples the composition appears to be consistent despite the DNA isolation kit that was used. As such, the overall microbial composition was not significantly different based on the DNA isolation protocol (PERMANOVA p = 0.87), with the exception of Sample 7, which displays high variability in both the relative abundance and NMDS plots. As expected, recovered microbes differed significantly across the 11 urine samples (PERMANOVA p = 0.001).

**Figure 3.**
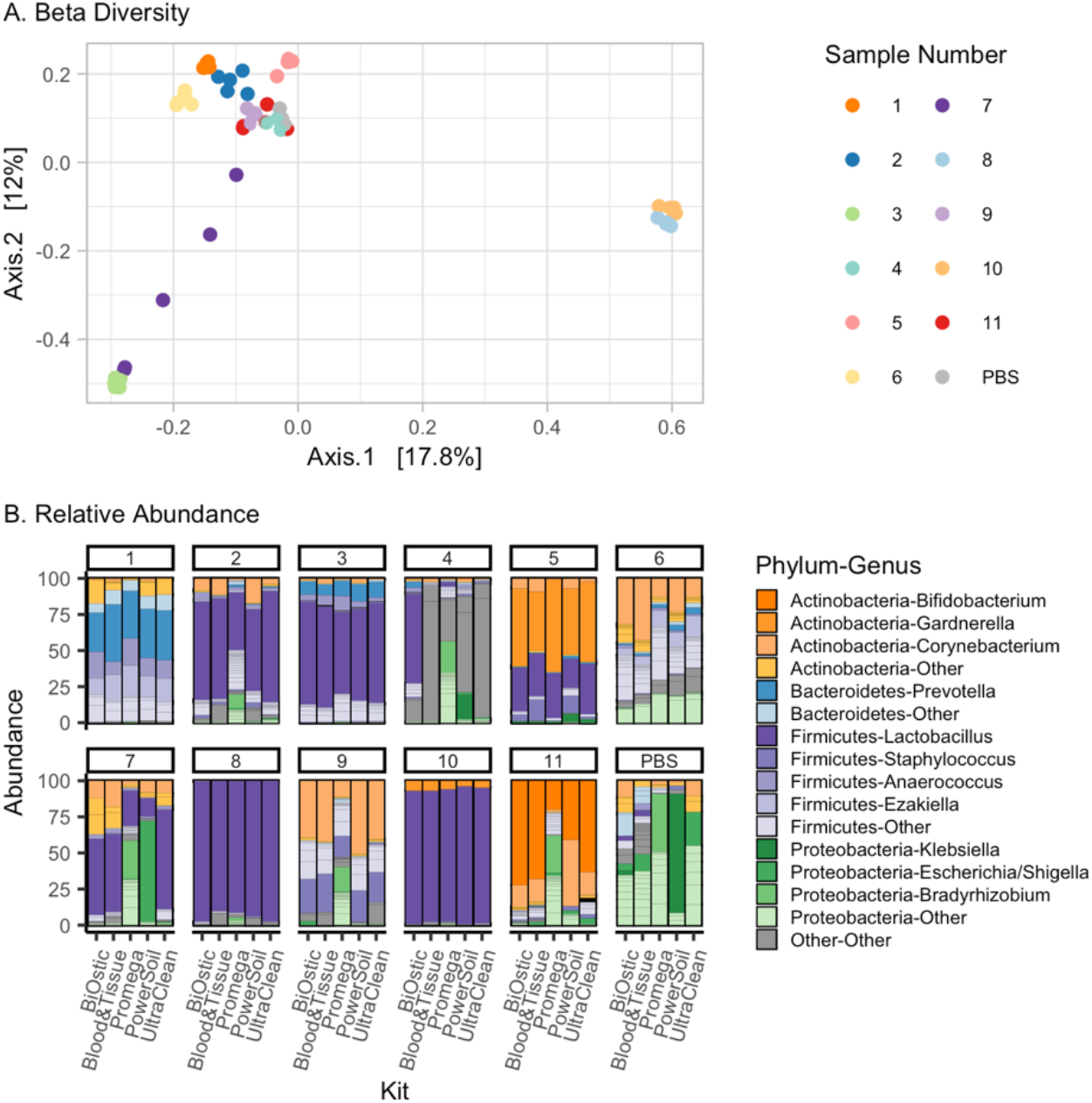
Differences in the DNA isolation methods do not result in drastic changes in relative abundances of identified genera. **A.** Multidimensional scaling plot using Bray Curtis distance demonstrates that most samples are not significantly different due to DNA isolation kit (p = 0.87 in PERMANOVA analysis), though samples 4 and 7 are not tightly clustered, indicating that these samples may have significant variations in microbiome composition by kit. **B.** Stacked bar plots represent the microbial composition of each sample after DNA isolation and 16S rRNA gene sequencing. Only sample 4, 7, and negative control PBS exhibit more variability.

### Recovery of Gram positive versus Gram negative bacteria

Prior studies comparing methods of DNA isolation from non-urine microbiome samples strongly indicated that the envelope structure of Gram positive organisms represents an impediment for uniform cell lysis. Therefore, we analyzed whether DNA isolation kits biased the identified microbial composition towards Gram negative species. We compared relative abundances among all genera with known Gram staining of representatives (Figure 4 and Figure S2 for individual sample results). Four out of five DNA isolation kits yielded comparable overall relative abundances of Gram positive bacteria. The Promega kit resulted in fewer Gram positive bacteria, though this was not statistically significant (Kruskal-Wallis, p = 0.197), likely due to the small sample size and highly variable data.

**Figure 4.**
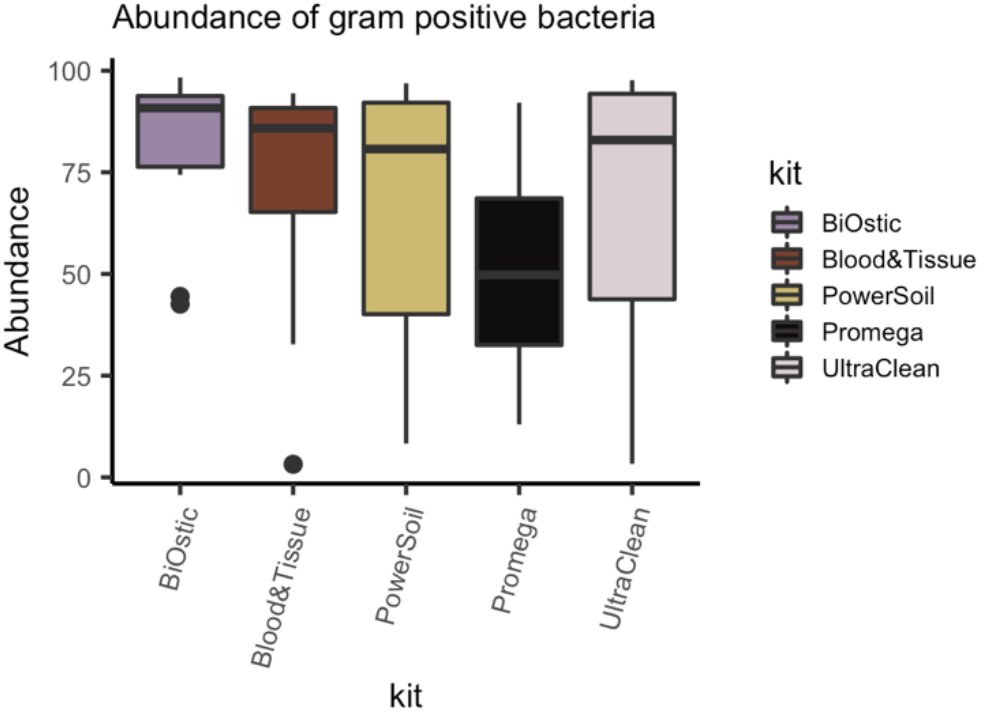
DNA from Gram positive bacteria was consistently presented in four out of five tested DNA isolation kits. The Promega kit recovered Gram positive bacteria, but at lower abundance than the other kits, thought this was not significantly different (Kruskal-Wallis, p = 0.197).

Microbiome studies of different niches reveal that overall composition is important for health and disease. However, infectious disease studies also show that specific microbes may be important for an underlying condition. Therefore, we further analyzed the presence of eight specific genera relevant for the urinary niche (Figure 5). These include genera containing three known urinary pathogens (*Escherichia, Klebsiella, Enterococcus*) and five genera typically considered as commensals (*Lactobacillus, Corynebacterium, Prevotella, Staphylococcus, Gardnerella*). Results from this analysis confirm our observation that the Promega kit is less efficient in extracting Gram positive bacteria, such as those belonging to the *Enterococcus, Corynebacterium,* and *Staphylococcus* genera. On the other hand, the Qiagen DNeasy PowerSoil kit appears to recover more of the ‘easy-to-lyse’ Gram negative organisms such as *Klebsiella* and *Escherichia* compared to the other kits.

**Figure 5.**
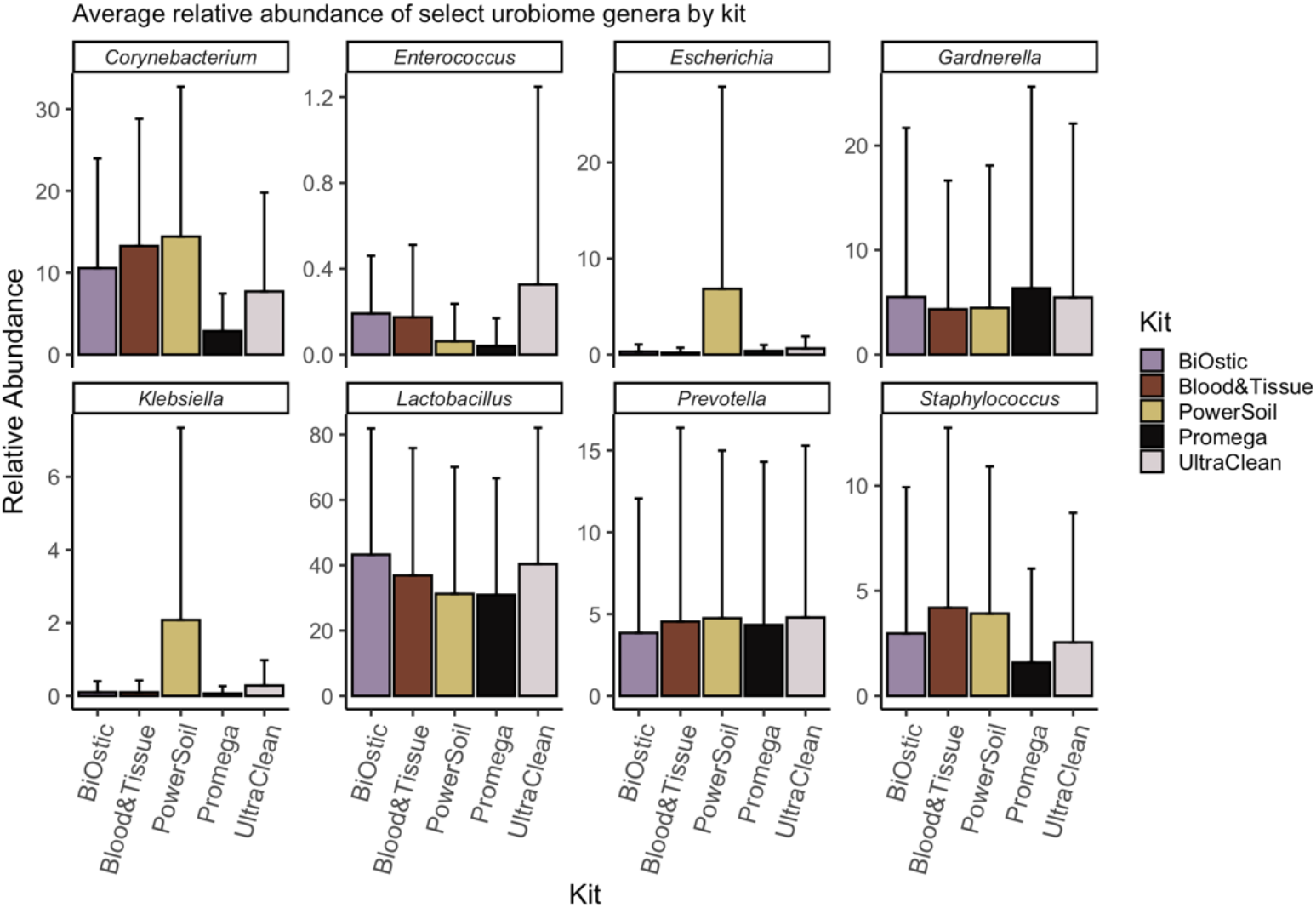
Comparison of relative abundances of genera with biologically-significant representatives. We compared relative abundances of bacteria recovered from eight genera with high biologic relevance including urinary pathogens (*Escherichia, Klebsiella, Enterococcus*) and commensals (*Lactobacillus, Corynebacterium, Prevotella, Staphylococcus, Gardnerella*). *Corynebacterium, Enterococcus, Lactobacillus,* and *Staphylococcus* are Gram positive bacteria and thus have cell walls that are more difficult to lyse during DNA isolation. *Gardnerella* are considered Gram variable while the *Escherichia, Klebsiella,* and *Prevotella* are considered Gram negative bacteria. The Promega kit tends to recover fewer Gram positive *Corynebacteria, Enterococci,* and *Staphylococci* compared to other kits while the PowerSoil kit recovers more *Escherichia* and *Klebsiella* compared to other kits.

## Discussion

The field of urinary microbiome research is still relatively new. As such, studies benchmarking DNA isolation kits and their performance in recovering urinary microbial composition data are lacking. This study aimed to compare several methods of isolating microbial DNA from human urine. In particular, we compared five commercially available DNA isolation kits and estimated not only the quantity of DNA, but also the quality of DNA when utilized in downstream compositional analyses.

It has previously been shown that biases are introduced to microbial composition analysis based on the DNA isolation technique in both high biomass and low biomass communities of microbes ^19,22,26^. Our results echo those found in oral microbial communities, where the DNA isolation method may result in significantly different DNA yield, though overall non-significant differences in downstream sequencing^23^. Though many of our downstream assessments showed non-significant differences, our data do not support the assumption that all DNA isolation kits perform equally in urinary microbiome studies, as we identified some important qualitative differences in recovery of Gram positive versus Gram negative organisms. Since microbiome data are presented in terms of relative abundance, if one type of microbe is absent due to a technical bias, it will artificially make other microbes appear more abundant. This is evident when viewing graphs in Figure S2, where relative abundances of Gram positive and Gram negative bacteria are inversely proportional to each other.

In our study, after initial PCR amplification, four samples and three controls derived from extremely low quantities of DNA failed to show a band on electrophoresis. The lack of amplified DNA after beginning with extremely low quantities of DNA could be expected. However, in one instance a sample extracted with the UltraClean kit had normal quantities of starting DNA with no evident PCR product on electrophoresis. This one result could have been spurious or possibly indicative of the presence of PCR inhibitors in the sample, as have been identified in other studies.

Our findings are strengthened by the multiple ways in which we assessed quality of DNA after isolation. This included evaluation of PCR products, assessment of the number of sequencing reads after high-throughput sequencing, as well as detailed compositional analyses of microbial data. We utilized an updated and rigorous bioinformatics pipeline to identify the genera corresponding to recovered sequences. We then utilized this information to assess the quality of sequencing information, which revealed important differences based on Gram staining characteristics and in urogenital genera that are highly relevant to the urinary microbiome field.

Our study certainly has multiple limitations, which are mainly related to technical factors. After assessing recovered DNA quantity using Qubit, we did not perform additional testing to assess the proportion of microbial versus human DNA contained in each sample. Thus, it is unclear if differences identified in total DNA recovery actually translate to differences in microbial DNA within different samples, which is the component of interest in microbiome studies. Another limitation was inherent in the need to divide urine samples. Though one urine sample was produced, we needed to ensure that it was equally divided prior to performing parallel testing with five kits. Since the biomass (e.g. cellular material containing DNA) may not be evenly distributed within the fluid of a urine sample, we addressed this issue by first centrifuging whole urine to produce a cell pellet containing the biomass. This cell pellet was then reconstituted in a smaller volume, thoroughly mixed, and then divided into five aliquots. However, it is still possible that due to pipetting or mixing errors, slightly different amounts of starting material were present in aliquots, which could have contributed to some of the variability seen in our results. However, we believe this factor is less important since urine volume did not correlate with biomass. For example, as shown in Tables S2 & S3, a 50mL sample (Sample 3) had the highest amount of recovered DNA while another 100mL sample had the lowest amount of recovered DNA. We utilized a negative control (PBS buffer) that was processed and sequenced in parallel to the urine samples. Though there was no starting added DNA in this sample, we recovered a small number of sequences (Figure S1) suggesting presence of low level contaminants. Unfortunately, we did not use separate controls at each analytic step and thus we are unable to distinguish the sources of the observed contamination, which could come from plastics in the laboratory, reagents within the DNA isolation kits, or during multiple technical steps prior to sequencing.

This study utilized voided urine, which is more reflective of the urogenital microbiome than the bladder microbiome. Since we are not attempting to characterize a niche, the method of urine sample acquisition is less important. However, microbes from the vagina are found in higher abundance in voided compared to catheterized urinary samples, and thus may have higher representation in the compositional data presented here. Since vaginal and urinary microbes are highly related in terms of the genera and species represented, vaginal contamination theoretically should not negatively impact the results of this benchmarking study^28,29^. Nevertheless, studies such as this one would ideally be replicated numerous times to confirm the findings.

## Conclusions

When considering the totality of our findings, DNA extracted with the Qiagen Biostic Bacteremia and DNeasy Blood & Tissue kits showed the fewest technical issues in downstream analyses, with the DNeasy Blood & Tissue kit also demonstrating the highest DNA yield. All five kits provided good quality DNA for high throughput sequencing with non-significant differences in the number of reads recovered, alpha, or beta diversity. However, in qualitatively assessing the types of bacteria, the Promega kit recovered fewer Gram positive bacteria compared to other kits.

The Promega and DNease PowerSoil kit also appear to have some important biases towards over-representing certain Gram negative bacteria of biologic relevance within the urinary microbiome. These findings have implications for research teams wishing to maximize utility of low biomass samples, particularly for sequencing strategies where more DNA is required. Furthermore, these findings are relevant for interpretation of microbiome studies. The results presented here are certainly in line with other microbiome niches suggesting that the DNA isolation methods used could potentially bias downstream results. As such, we urge caution to investigators when selecting which DNA isolation method is used in future urinary microbiome studies, caution to the scientific community when assessing findings from studies where isolation methods with known bias were used, and further urge a high level of caution in general when trying to compare or extrapolate results from studies where different DNA isolation methodologies were used.

## Materials and Methods

### Sample collection and processing

This study was deemed exempt by the Duke University Institutional Review Board (Pro00085111). Following all relevant guidelines, de-identified voided urine samples were collected in sterile cups from the Duke Urogynecology clinic, refrigerated (4°C), and processed within 4-10 hours (Table S2). As the study was deemed exempt by IRB no consent was obtained. During processing, samples were handled aseptically, transferred to 50 mL conical tubes and spun to collect all of the biomass, including human and microbial cells (4°C, Eppendorf 5810R centrifuge, 15 min, 3,220 rcf) represented in the “cell pellet”. Supernatants were decanted and the remaining cell pellets with residual urine were transferred into sterile 1.5 mL tubes, then spun again at 10,000 rcf in the Eppendorf 5340R centrifuge for 5 min at 4°C. The total cell pellet per sample was resuspended in sterile filtered phosphate buffered saline (PBS) on ice. Re-suspended pellets were divided into 5 identical aliquots, and stored at −80°C until DNA isolation.

### DNA isolation procedures

This step started with the five identical aliquots and thus the same starting material was processed in parallel with five commercially available DNA isolation kits. Each kit had differing levels of chemical, mechanical, and enzymatic cell lysis, as summarized in Table 1. PBS buffer was used as a negative control sample with each DNA isolation kit. For the Qiagen DNeasy Blood & Tissue kit we performed the optional steps as recommended in the protocol for optimizing recovery of gram-positive bacteria. All samples were assessed using the Agilent 2100 Bioanalyzer, Promega GlowMax spectrophotometer and ThermoFisher Qubit HR reagents to determine the quality and quantity of recovered DNA. Recovered DNA concentrations are provided in Table S3.

### Bacterial ribosomal DNA amplification and sequencing

DNA samples and negative control were subjected to PCR in order to amplify the V4 variable region of the 16S rRNA gene. For PCR, forward primer 515 and reverse primer 806 were used following the Earth Microbiome Project protocol (http://www.earthmicrobiome.org/). These primers (515F and 806R) carry unique barcodes allowing for construction of a library of pooled samples for sequencing. PCR products were quantified and pooled. In instances where no PCR product was detected (see Table S3), equivalent volumes of the final PCR amplification solution were pooled with the others. Combined pooled samples were then submitted for sequencing on an Illumina MiSeq sequencer configured for 150 base-pair paired-end sequencing runs. DNA samples for all kits were prepared and sequenced together to avoid processing and sequencing batch variations.

### Sequencing data processing and analysis

Raw sequences were trimmed and de-multiplexed prior to being processed with DADA2 (v.1.14.0) to provide amplicon sequence variants (ASVs) per sample^30^. ASVs were compared against the SILVA reference database (v.132) using the RDP classifier implemented in DADA2 for identification of taxa prior to being analyzed with phyloseq and vegan in R^31–33^. Overall microbial composition was assessed by estimating alpha diversity (number of observed genera, Shannon Index, and Inverse Simpson Index) as well as beta diversity using the Bray-Curtis distance. Comparisons across the 5 isolation kits were statistically evaluated using the Kruskal-Wallis rank sum tests followed by pair-wise comparisons using Wilcoxon-rank sum tests, and PERMANOVA for Bray-Curtis distances.

## Data Availability

All sequences are available for download in the Sequence Read Archive under Accession Number PRJNA PRJNA662669 (http://www.ncbi.nlm.nih.gov/bioproject/662669).

## Acknowledgements

We thank the clinical research coordinators (Shantae McLean, M.P.H., Robin Gilliam, M.S.W., Akira Hayes, M.S.) at the Duke Urogynecology Clinic who assisted in collecting urine samples for this study. We also thank the Duke University School of Medicine for the use of the Microbiome Shared Resource (in particular to Holly Dressman, Ph.D.), which provided Promega kit isolation and all the library preparation services, as well as the Sequencing and Genomic Technologies Core Facility of the Duke University Center for Genomic and Computational Biology. This study was funded by: K12 Duke KURe (DK100024) NIDDK and UAH Startup fund to TAS; career development awards K23 DK110417, R03 AG060082, and P30 AG028716 to NYS; and K01 DK116706 to LK.

## Author Contributions

TAS conceived the study and performed the experiments; TZ and AB assisted with literature review, analysis and data annotation; TAS, LK, and NYS analyzed the data; CLA contributed in discussing the results and editing the manuscript. TAS, LK and NYS interpreted the data and wrote the manuscript. All authors read and approved the final manuscript.

