## Supplemental Materials for "Benchmarking DNA isolation kits used in analyses of the urinary microbiome"

**Table S1. Diverse DNA isolation methods utilized in analyses of microbial composition of lower urinary tract.**

| Study | Urine Collection Method | Volume | DNA Extraction Kit | Lysis Method | DNA Collection Method |
|---|---|---|---|---|---|
| Nelson 2010 | Void | <50mL | DNeasy Blood and Tissue Kit | Enzymatic | Silica spin column |
| Dong 2011 | Urethral swab or void | 1mL or 5mL | DNeasy Tissue Extraction Kit | Enzymatic | Silica spin column |
| Siddiqui 2011 | Void | 30mL | DNeasy Blood and Tissue Kit | Enzymatic | Silica spin column |
| Fouts 2012 | Void or catheterization | -- | Custom with lysozyme/bead lysis and phenol extraction | Enzymatic and bead beating | Ethanol precipitation |
| Wolfe 2012 | Void, aspiration, and catheterization | -- | DNeasy Tissue Extraction Kit | Enzymatic | Silica spin column |
| Lewis 2013 | Void | 2mL | Custom with SDS/bead lysis and alcohol precipitation | Bead beating | Isopropanol and ethanol precipitation |
| Hilt 2014 | Catheterization | 1mL | DNeasy Blood and Tissue Kit | Enzymatic | Silica spin column |
| Pearce 2015 | Catheterization | 1mL | HMP (Yuan 2012) and DNeasy Blood and Tissue Kit | Enzymatic | Silica spin column |
| Shoskes 2016 | Void | -- | PowerMag Microbiome RNA/DNA Isolation Kit | Bead beating | Magnetic beads |
| Karstens 2016 | Catheterization | 50mL | DNeasy Blood and Tissue Kit | Enzymatic | Silica spin column |
| Modena 2017 | Void | 50mL | Custom with TRIzol | TRIzol Reagent | Alcohol precipitation |
| Thomas-White 2017 | Void or catheterization | 1mL | DNeasy Blood and Tissue Kit | Enzymatic | Silica spin column |

| | | | | | |
|---|---|---|---|---|---|
| Gottschick 2017 | Void | 15mL | Phenol extraction with consecutive peqGOLD Tissue DNA Kit | Bead beating | Silica spin column |
| Liu 2017 | Modified void | 40mL | Custom with PowerMag Microbiome RNA/DNA Isolation Kit | Bead beating | Magnetic beads |
| Popovic 2017 | Void | 30mL | PowerSoil DNA Isolation Kit | Bead beating | Silica spin column |
| Wu 2017 | Catheterization | 50mL | DNeasy Blood and Tissue Kit | Enzymatic | Silica spin column |
| Burton 2017 (dog urine) | Aspiration (cystocentesis) | 30mL | Custom with QIAamp | Bead beating | Silica spin column |
| Adebayo 2017 | Void | 1mL | DNeasy Blood and Tissue Kit | Enzymatic | Silica spin column |
| Shrestha 2018 | -- | 30mL | Custom with phenol extraction | Enzymatic and bead beating | Alcohol precipitation |
| Jung 2019 | Void | 200µL | DNeasy Powersoil Kit | Bead beating | Silica spin column |
| Pohl 2020 | Void and catheterization | -- | DNeasy or QIAmp DNA Micro Kit | Enzymatic | Silica spin column |
| Forster 2020 | Catheterization | >5mL | QIAamp Circulating Nucleic Acid Kit with QIAamp columns | Enzymatic | Silica column |
| Kinneman 2020 | Catheterization | >1mL | EZ1 DSP Virus Kit | Enzymatic | Silica magnetic beads |

**Table S2. Approximate volumes* of the void urine used in this study.**

| Sample | Volume |
|---|---|
| 1 | 30 mL |
| 2 | 30 mL |
| 3 | 50 mL |
| 4 | 75 mL |
| 5 | 75 mL |
| 6 | 50 mL |
| 7 | 75 mL |
| 8 | 75 mL |
| 9 | 100 mL |
| 10 | 100 mL |
| 11 | 100 mL |

**Volume estimates were measured in the cups provided, without additional transfer into measuring cylinders to avoid contamination and therefore, these are estimates that are rounded to closest measuring mark.*

**Table S3. Recovered concentrations after DNA isolation**

| Sample/kit | Kit 1 (BiOstic) | Kit 2 (Blood&Tissue) | Kit 3 (Promega) | Kit 4 (PowerSoil) | Kit 5 (UltraClean) |
|---|---|---|---|---|---|
| 1 | 0.255 | 1.107 | 0.000 | 0.012 | 0.032 |
| 2 | 0.282 | 0.584 | 0.000 | 0.126 | 0.101 |
| 3 | 4.763 | 9.402 | 0.578 | 4.563 | 1.576 |
| 4 | 0.995 | 2.386 | 0.066 | 0.808 | 0.695* |
| 5 | 2.046 | 4.403 | 0.063 | 1.138 | 0.001 |
| 6 | 0.101 | 0.395 | 0.000 | 0.040 | 0.037 |
| 7 | 0.804 | 1.027 | 0.000 | 0.137 | 0.269 |
| 8 | 1.153 | 5.752 | 0.121 | 0.906 | 0.751 |
| 9 | 0.549 | 1.333 | 0.078 | 0.265 | 0.331 |
| 10 | 1.362 | 3.073 | 0.240 | 0.745 | 0.596 |
| 11 | 0.007* | 0.055 | 0.000 | 0.004* | 0.000* |
| PBS control | 0.000* | 0.008 | 0.000 | 0.005* | 0.007* |

*DNA concentrations in ng/μL*

** - no agarose gel band detected after PCR amplification*

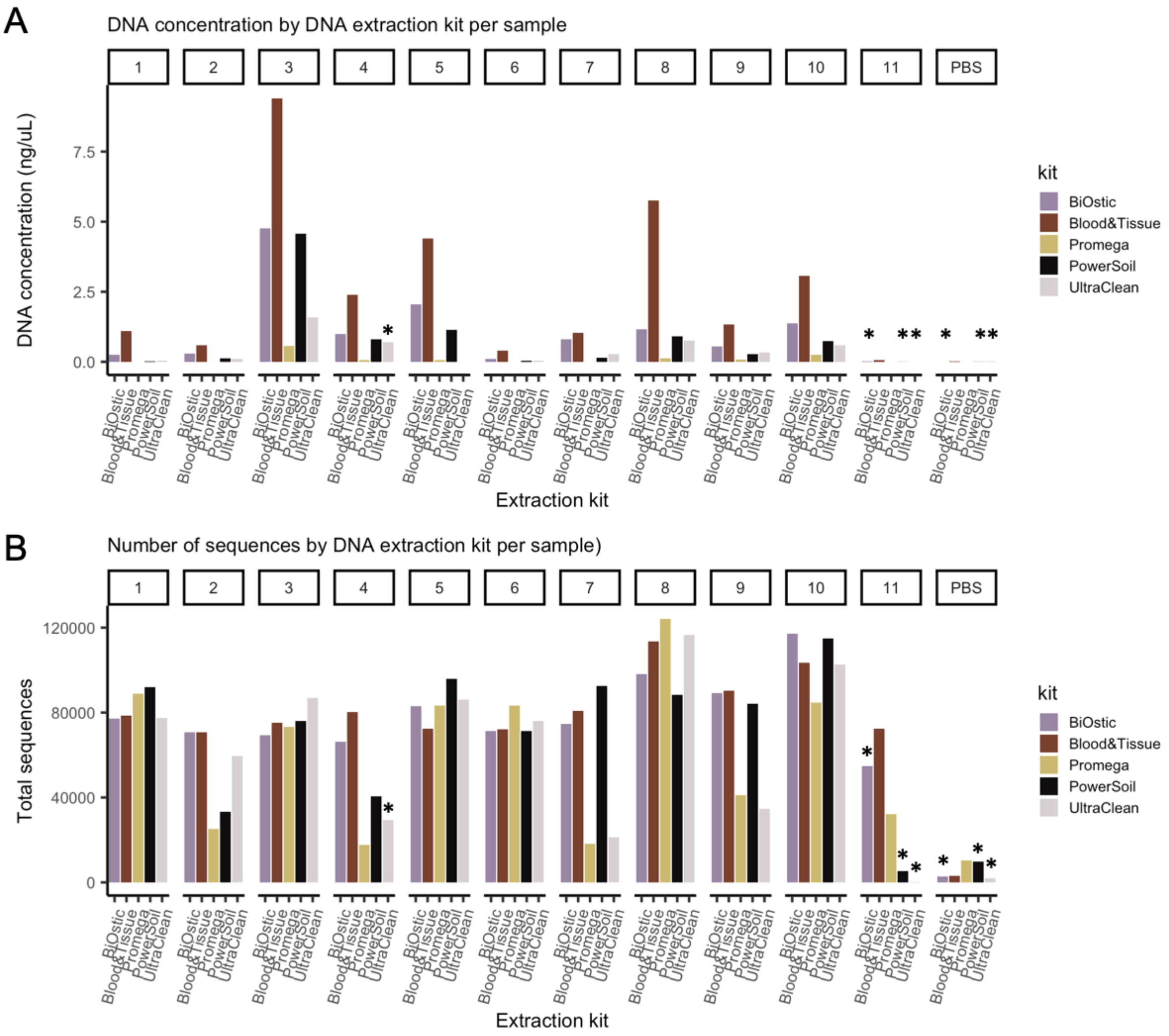


**Figure S1. Comparative data for all samples. A.** Comparing the amount of isolated DNA from urine. Asterisks indicate samples that did not produce identifiable PCR product. **B.** Comparing the number of sequencing reads obtained from DNA isolated with each kit. Samples 1 - 11 represent DNA extracted from voided urine while sample #12 is a negative control of filtered sterile PBS solution. Asterisks in both panels mark samples that did not yield detected gel bands after amplification of isolated DNA with primers specific for the V4 region of the 16S rRNA gene.

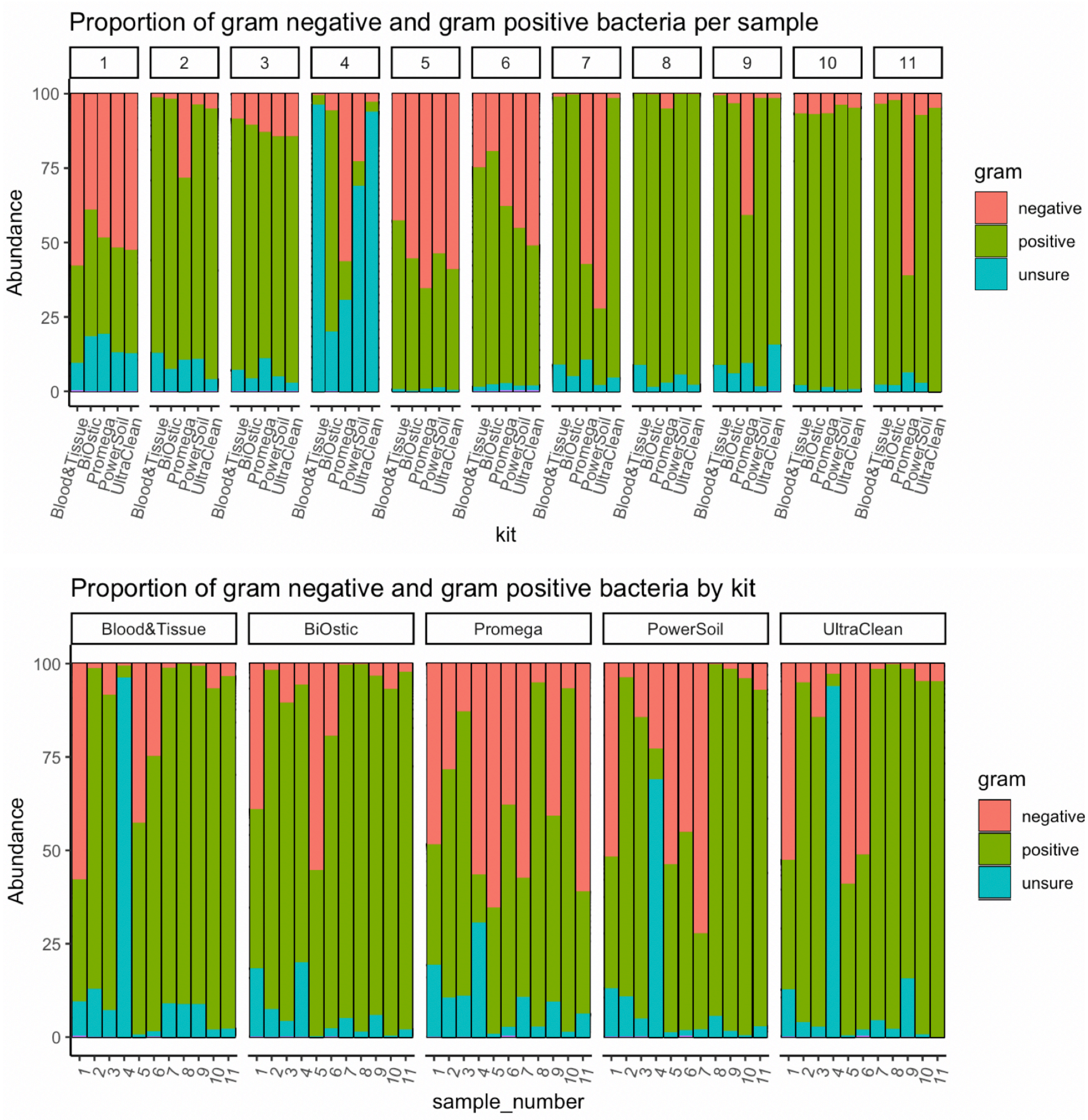


**Figure S2. Relative abundance of Gram positive and Gram negative bacteria in per sample or per kit basis.** The summary of these data is presented in Figure 5.
